# Small-spored *Alternaria* spp. (section *Alternaria*) are common pathogens on wild tomato species

**DOI:** 10.1101/2022.12.08.519636

**Authors:** Tamara Schmey, Corinn Small, Lina Muñoz Hoyoz, Tahir Ali, Soledad Gamboa, Betty Mamami, German C. Sepulveda, Marco Thines, Remco Stam

## Abstract

The wild relatives of modern tomato crops are native to South America. These plants occur in habitats as different as the Andes and the Atacama Desert and are to some degree all susceptible to fungal pathogens of the genus *Alternaria. Alternaria* is a large genus. On tomato, several species cause early blight, leaf spot, and other diseases.

We collected *Alternaria*-like infection lesions from the leaves of eight wild tomato species from Chile and Peru. Using molecular barcoding markers, we characterized the pathogens. The infection lesions were caused predominantly by small-spored species of *Alternaria* of the section *Alternaria*, like *A. alternata*, but also by *Stemphylium* spp., *Alternaria* spp. from the section *Ulocladioides*, and other related species. Morphological observations and an infection assay confirmed this. Comparative genetic diversity analyses show a larger diversity in this wild system than in studies of cultivated *Solanum* species.

As *A. alternata* has been reported to be an increasing problem on cultivated tomato, investigating the evolutionary potential of this pathogen is not only interesting to scientists studying wild plant-pathosystems. It could also inform crop protection and breeding programs to be aware of potential epidemics caused by species still confined to South America.

## Introduction

Host-pathogen interactions are determined by environmental conditions that can determine the level of success to which a pathogen can proliferate and cause disease on different host genotypes (Agrios, 2005). Surprisingly, there have been limited investigations into wild pathosystems, with few notable exceptions. The main pathosystem supporting the gene-for-gene model is the well-studied flax (*Linum*) - flax rust (*Melamspora*) association. Local adaptations were quantified showing that more virulent *Melamspora* strains occurred predominantly on more resistant flax, conversely susceptible flax harbored avirulent flax rust strains suggesting continuous selection pressures occurring between host and pathogen (Thrall *et al*., 2002; Thrall and Burdon, 2003). Investigations into the wild *Plantago lanceolata*-Powdery mildew (*Podosphaera plantaginis*) system found variation in infectivity depending on the degree of host sympatry and described spatiotemporal variation of infectivity giving rise to regional heterogeneity (Laine, 2005; Soubeyrand *et al*., 2009). Co-evolutionary forces have been observed in *Dianthus* - anther smut (*Microbotryum*) associations, where co-occurrence (competition in a single host population) was observed between host-specialized differentiated *Microbotryum* lineages (Petit *et al*., 2017). Sea rocket *(Cakile maritima)*, a coastal crucifer species, experiences regular, but short-lived epidemics of *Alternaria brassisicola* that vary based of age of the plants, as well as location and climatic conditions (Thrall *et al*., 2001). Although some of these wild systems are well-studied, overall literature on wild plant pathogens remains scarce.

Wild tomato species are interesting model species. They have speciated both sympatrically and allopatrically due to low gene flow and genetic drift between populations and are diverse within and between populations. As relatives to the cultivated tomato *S. lycopersicum*, they have been frequently sampled, sequenced, and have been used in ample phylogenetic and genomic analyses (Städler *et al*., 2005, 2008; Peralta *et al*., 2007; The 100 Tomato Genome Sequencing Consortium *et al*., 2014; Stam *et al*., 2019; Alonge *et al*., 2020). As useful models for studying diversity in resistance genes, wild tomatoes also provide a great potential to understand the relationship of Resistance-gene diversity and diversity among pathogens (Dodsworth *et al*., 2016).

In particular, the species *Solanum chilense* has been widely studied in the context of climatic adaptation and pathogen resistance. This plant species originated in southern Peru, migrated south towards northern Chile (Städler *et al*., 2008; Böndel *et al*., 2015; Beddows *et al*., 2017) and is now composed of several genetic subgroups. The two southern lineages that migrated along the coast and into the mountains are genetically more similar to the central group than to each other (Böndel *et al*., 2015; Stam *et al*., 2019). As such, *S. chilense* occurs in a wide range of habitats and exhibits local adaptations to both abiotic and biotic factors (Peralta *et al*., 2008; Chetelat *et al*., 2009; Fischer *et al*., 2013; Böndel *et al*., 2015; Nosenko *et al*., 2016; Stam *et al*., 2017, 2019). The species also shows variation in resistance towards economically important pathogens including *Phytophthora infestans, Alternaria solani, Fusarium species* and *Cladosporium fulvum* (syn. *Passalora fulva)* (Stam *et al*., 2017, 2019; Kahlon *et al*., 2020). Additionally, significant genetic diversity exists among resistance genes (Stam *et al*., 2019; Kahlon *et al*., 2020) and populations show clear variation in quantitative defense responses within an between populations (Kahlon *et al*., 2021). Specific horizontal or vertical resistance properties against a range of pathogens have been identified in other tomato spp. as well; certain *S. habrochaites* accessions possess vertical (qualitative) resistance against the bacterial spec disease *Pseudomonas syringae* (Hassan *et al*., 2017), as well as horizontal (quantitative) resistance against *P. infestans* (Li *et al*., 2011). Several wild tomato species show differences in resistance against *P. fulva* (Kruijt *et al*., 2005), for example *S. pimpinellifolium* displays a clear north-south gradient for horizonal (Resistance gene-mediated) resistance against this pathogen (Van der Hoorn *et al*., 2001).

For this study, we sampled on eight wild tomato species, seven of which belong to *Solanum* section *Lycopersicon. Solanum lycopersicoides* belongs to the section *Sitiens* and is therefore more distantly related to the other species, which can already be seen from a divergent flower morphology and its characteristic black fruits. The section *Lycopersicon* comprises four subsections (Pease *et al*., 2016), which are all represented by our host species: *S. habrochaites* and *S. pennellii* represent subsection *Hirsutum*; *S. peruvianum, S. corneliomulleri* and *S. chilense* represent subsection *Peruvianum* and *S. arcanum* represents subsection *Arcanum*. These species share a complex relationship but have different habitats and features that can be visually distinguished in the field (Peralta *et al*., 2008; Dodsworth *et al*., 2016; Knapp and Peralta, 2016; Beddows *et al*., 2017). The subsection *Esculentum* is represented by *S. pimpinellifolium*. It is the only red-fruited species in this study and potentially the wild progenitor of the cultivated tomato *S. lycopersicum* (Knapp and Peralta, 2016; Razifard *et al*., 2020). Species ranges and habitats vary for all eight host species and cover various conditions from temperate and high-altitude environments to arid regions on the borders of the Atacama (Peralta *et al*., 2008).

Cultivated tomatoes are hosts to many pathogens. Two of the most dominant leaf pathogens are the causal agents for early blight or leaf spots and late blight, *Alternaria* spp. and *Phytophthora infestans*, respectively. Lindqvist-Kreuze et al. 2019 have identified *P. Infestans* on several wild tomato plants in high altitude habitats, in the vicinity of potato fields (Lindqvist-Kreuze *et al*., 2020). Recently we sampled the phyllosphere microbiome of four distinct wild tomato species in two regions in Peru. Specifically targeting leaves with infection symptoms throughout these regions, we found that *Alternaria* spp. are omnipresent on the leaves, but that *P. Infestans* was rare or absent (Runge *et al*., 2022). Taking into account that we and other studies previously found clearly quantifiable differences in resistance against *A. solani* between and within wild tomato populations (Chaerani and Voorrips, 2006; Stam *et al*., 2017) we focused specifically on the diversity of *Alternaria* and closely related fungal genera.

These genera are part of the phylum *Ascomycota* (family *Pleosporaceae*) and are composed of species living a wide variety of lifestyles (Woudenberg *et al*., 2013). Small-spored *Alternaria* from section *Alternaria*, like *A. alternata*, can cause leaf spot and other diseases on a plethora of hosts (Woudenberg *et al*., 2015). *A. solani* is more host-specific to the nightshade (*Solanaceae*) family, occurring commonly on domesticated tomato (Song *et al*., 2011; Kumar *et al*., 2013). Early blight on tomato can be caused by *A. solani* as well *as A. linariae*, which was previously called *A. tomatophila* (Adhikari *et al*., 2020). The three species *A. alternata, A. solani* and *A. linariae* can cause similar lesions in potato and tomato (Adhikari *et al*., 2020). The first symptoms are necrotic lesions, which are small and dark. Larger early blight lesions become target-like with concentric rings and a yellowing zone around the lesion (Chaerani and Voorrips, 2006). Woudenberg *et al*. 2013 argue that the genus *Ulocladium* is synonymous to *Alternaria*, as phylogenetic analysis places several *Ulocladium* species within *Alternaria* (Woudenberg *et al*., 2013). *Ulocladium atrum*, now *Alternaria atra*, regularly infects *Solanum* spp. (Norse, 1974; Esfahani, 2018). *Stemphylium* is another *Pleosporaceae* genus which causes grey leaf spot on tomato. Grey leaf spot lesions are often smaller, but in some cases resemble *Alternaria*-caused symptoms. The disease is of relatively lower economic importance than early blight, but has been documented globally, including recently in the Venezuelan Andes (Cedeño and Carrero, 1997).

Recent studies have used different genetic markers to elucidate the diversity of *A. solani* and *A. alternata* found on domesticated potato and tomato crops (Ding *et al*., 2019; Adhikari *et al*., 2020). For the current study, we chose four barcode markers (Woudenberg *et al*., 2015) based on the discriminatory power and feasibility reasons, namely the internal transcribed spacer region ITS1F, the RNA polymerase second largest subunit (RPB2), the translation elongation factor 1-alpha (TEF1) and the *Alternaria* major allergen gene (Alt a 1).

Here we present the results of a targeted sampling strategy. We collected visually symptomatic leaves with necrotic lesions from wild tomato species in six ecologically diverse regions in Chile and Peru and extracted the causal fungi. Using barcode sequencing, we gained first insights into the diversity of these pathogens.

## Experimental procedures

### Sampling locations and collection

In February 2018 and 2019, we visited and sampled from a total of 81 sites near the following cities: Cajamarca, Tacna and Lima in Peru as well as Arica, Antofagasta and San Pedro de Atacama in Chile. Sampling sites included both previously visited sites (as documented by the Tomato Genetics Resource Center TGRC http://tgrc.ucdavis.edu) and newly discovered sites. The sites extended across various climatic zones and the environments consisted of coastal terrain, the high west Andean plains (1500-2500 m) and mountainous terrain. Environments in which tomato populations were found varied in temperature, precipitation, UV exposure and atmospheric oxygen concentrations. Biotic factors also varied (i.e., local plant species composition and densities). Major *Solanum chilense* areas were located in three previously defined regional subgroups spanning Chile and Peru (Böndel *et al*., 2015). These subgroups were defined as Central, Southern low-altitude and Southern high-altitude. Other *Solanum* species were sampled throughout these ranges and two additional regions farther north near Lima and Cajamarca (Fig. 1). The sole wild tomato species in two southernmost sampling regions is *S. chilense*. In the central regions (Tacna and Arica), *S. chilense, S. lycopersicoides* and *S. peruvianum* were found while *S. arcanum, S. habrochaites, S. corneliomulleri*, and *S. pimpinellifolium* occurred in the northernmost regions near Lima and Cajamarca.

**Figure 1:**
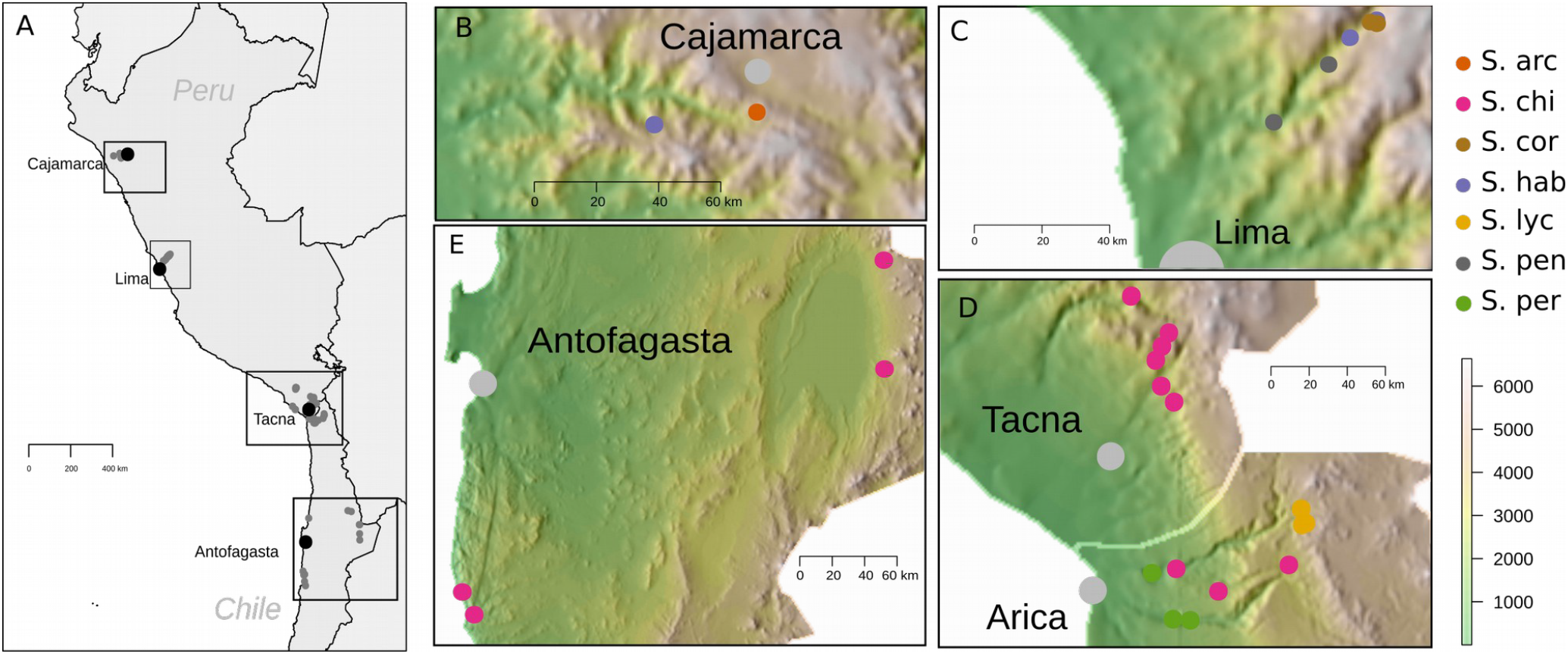
Map of sampling regions A) overview map of Peru and northern Chile, indicating all locations where samples were collected from wild tomatoes (grey dots). Major cities are indicated (black dots) and the four major geographic areas presented in B-E are boxed approximately. B-E) Maps of the four major geographic areas where *Alternaria*-related samples were collected. The dots represent sites from which samples have been isolated, purified and sequenced. Each dot is color coded to represent the dominant host species at the site (orange: *Solanum arcanum*, magenta: *S. chilense*, hazelnut: *S. corneliomulleri*, mauve: *S. habrochaites*, mustard: *S. lycopersicoides*, dark grey: *S. pennellii*, green: *S. peruvianum*.) Major cities are indicated with light grey circles. Panel E shows two sampling regions; coastal locations will be referred to as Antofagasta region or southern coast and eastern locations as the region around San Pedro de Atacama or southern highlands. Elevations are shown in m.a.s.l.

Population sizes varied among sites, ranging from one plant to several hundred individual plants. We packed the samples in paper, stored them in coolers during collection day trips and transferred them to a cool, dry storage until sample purification. We collected leaves displaying any blight-like symptoms from young and old plants. When we observed typical *Alternaria* Early Blight-like symptoms (circular, ringed, “bullseye” lesions) we prioritized these leaves for collection (Fig. 2).

**Figure 2:**
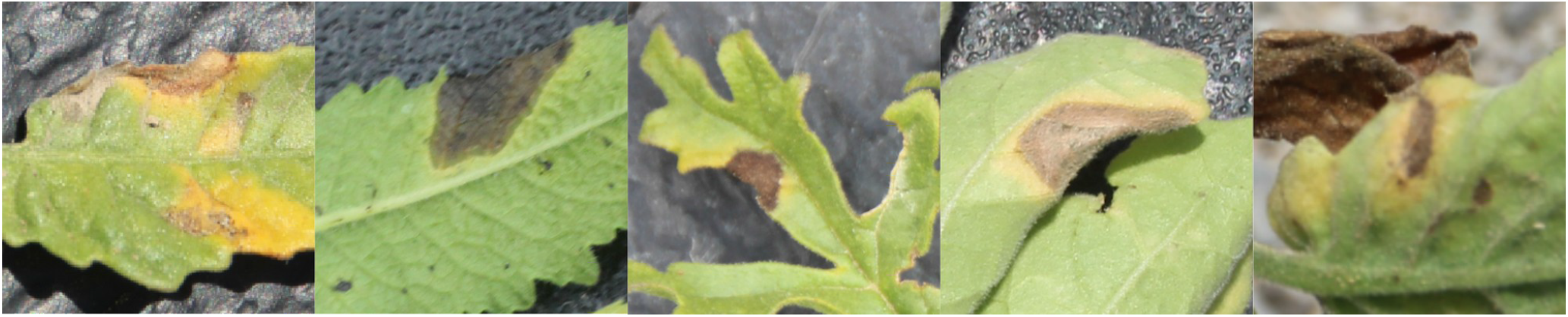
Infection lesions Each panel represents typical disease symptoms as observed on different hosts in the field. From left to right: *S. chilense, S. habrochaites, S. lycopersicoides, S. peruvianum, S. pimpinellifolium*.

### Sample purification

Once the leaves were dried, we submerged them in 2% bleach for 3 minutes to sterilize the leaf surface, subsequently washed them with water for 3 minutes and plated them on synthetic nutrient-poor agar (SNA) plates (7.34 mM KH2PO4, 9.89 mM KNO3, 2.03 mM MgSO4-7H20, 6.71 mM KCl, 1.11 mM glucose, 0.58 mM saccharose, 65.41 mM agar, 0.6 mM NaOH) to promote *Alternaria* spp. growth. Plates with initial leaf samples incubated at 15 °C and 8 hours of UV light per day. To minimize contamination, we monitored for various types of fungal growth each day. From all fungi resembling *Alternaria* and related species, we prepared subcultures with tiny agar plugs from clean growth and transferred them to fresh SNA plates. All subcultures grew under the above-mentioned conditions until they showed sufficient visible growth. Furthermore, we generated subcultures on SNA plates with pieces of sterilized filter paper for cryopreservation. When spores were visible, we transferred the filter paper to cryo-tubes which we froze using liquid nitrogen and stored at -80 °C. To visually identify species, we monitored the fungal growth for spores. We examined both hyphal structure and spore morphology under a binocular stereo-microscope and a compound microscope, respectively.

### DNA extraction and molecular characterization

For DNA extraction, we grew mycelium in liquid culture. We transferred agar plugs of fresh growth from clean cultures to Erlenmeyer flasks containing 100-200 ml of autoclaved Potato Dextrose Broth (PDB, Roth, Karlsruhe, Germany) and incubated at 24-28 °C shaking at 100 rpm exposed to light. Following 4-7 days of incubation, most flasks had a sufficient amount of mycelial growth without significant melanization (which hampers DNA extraction). To collect the mycelium, we filtered the liquid culture through sterilized grade 90 cheesecloth, washed the flask with MQ water and squeezed the collected mycelia to remove liquid. After transferring the mycelia to 15 or 50 ml centrifuge tubes, we placed the tubes into liquid nitrogen and lyophilized overnight to remove all remaining liquid. We prepared a 96-well plate by adding smaller and larger magnetic beads in a ratio of 3:1 to each 2ml well. Then we placed 5mg dry-weight lyophilized tissue into the wells. The 96-well plates were sealed and shipped in dry ice overnight.

Using the KingFisher Flex (Thermo Fisher Scientific Inc., Waltham, MA) with the BioSprint DNA Plant kit (Qiagen, Hilden, Germany) we extracted and purified genomic DNA from the tissue samples following the manufacturers’ instructions. Then we amplified and sequenced four genomic barcode markers to molecularly characterize the samples. These four markers were the targeted Internal Transcriber Spacer 1 (ITS1F) rDNA, Alternaria major allergen (Alt a 1), RNA polymerase second largest subunit (RPB2) and translation elongation factor 1-alpha (TEF1). We carried out the PCR reactions in 25 μl [13.6 μl ddH2O, 5 μl MangoTaq reaction buffer colorless, 25 μl dNTPs (2 mM each), 1 μl MgCl2 (50 mM), 1 μl bovine serum albumin (BSA, 20 μg/μl), 0.4 μl forward primer (25 μM), 0.4 μl reverse primer (25 μM), 0.1 μl Mango Taq DNA polymerase and 1 μl DNA sample] with the primers listed in supplementary table 1 and PCR conditions as described in supplementary table 2. Additionally, we purified the resulting PCR product of ITS1F using the QIAquick Gel Extraction Kit (Qiagen, Hilden, Germany). Finally, we sequenced all PCR products on the ABI Prism 377 DNA Sequencer using BigDye (Applied Biosystems, Foster City, CA) Cycle Sequencing Kit version 3.1 with the forward primer from the PCR reaction.

### Phylogenetic analyses

Sequence data processing and phylogenetic analyses employed the in-house developed pipeline AB12PHYLO (Kaindl *et al*., 2022). With this pipeline, we checked the sequencing results for quality and relevance for the study. The AB12PHYLO pipeline performs a first assessment of the read quality. Only isolates that have sufficient sequence quality at all four loci are kept for phylogenetic analyses. The pipeline includes the possibility to obtain species identifications based on hits from BLAST searches in the NCBI database. We inspected these BLAST hits to check the relevance of the species.

To include reference species in the phylogenetic tree, we downloaded sequence data from the NCBI GenBank database (Accession numbers of reference sequences are listed in supplementary table 3). Then, we constructed a raw phylogenetic tree, from which we subsequently removed a clade of samples that grouped together with unrelated species like *Fusarium* spp. and *Cladosporium* spp. etc. This approach removes not only unrelated species but also isolates that group together with these species due to poor sequence quality. Visual inspection of branch lengths and the multiple sequence alignment (msa) confirmed the necessity to remove all isolates from this clade, even if the blast search gave a hit for *Alternaria*. Finally, we used the combined set of sequence data for the remaining 139 isolates plus the reference sequences to build a phylogenetic tree of concatenated markers with AB12PHYLO. The AB12PHYLO pipeline aligned the sequences using mafft (Katoh *et al*., 2002), trimmed the multiple sequence alignments in a balanced setting based on adjusted settings in gblocks (Castresana, 2000) and concatenated the trimmed alignments. Then it inferred the Maximum Likelihood (ML) tree using RAxML-NG (Kozlov *et al*., 2019) set to 20 random and 20 parsimony-based starting trees, 1000 bootstrapping iterations and evomodel: GTR+G. The node support values are Transfer Bootstrap Expectations (TBE) (Lemoine *et al*., 2018). The samples with blast hits for *Stemphylium* and *Pleospora* were closer to the reference for *Stemphylium vesicarium* than to the reference for *Stemphylium botryosum*. Consequently, we rooted the tree with *Stemphylium botryosum* as outgroup. Besides constructing a multigene phylogeny, we also used AB12PHYLO to construct trees for each of the phylogenetic markers separately with the same settings.

### Pathogen distribution and diversity

To provide an overview which pathogens were found on which host plants and in which regions, we visualized the distribution in an alluvial plot using the package ggalluvial in R. The AB12PHYLO pipeline has a functionality to calculate nucleotide diversity statistics from the sequence alignments. We used it to generate statistics for each of the pathogen groups in the concatenated tree. Furthermore, we calculated statistics for the different sampling regions.

### Morphological characterization of conidia of selected samples

For morphological characterization, we cultured 14 isolates on SNA plates at 25°C, 12 h UV-A light, 12 h darkness and 85% humidity for eight days. Using a scalpel, we scraped the spores from the plates and placed them in a drop of water on microscope slides. Subsequently, we took pictures of the spores under the microscope Axio Imager.Z1 (Brightfield) with the camera AxioCamHR (both from Carl Zeiss Microscopy Deutschland GmbH, Oberkochen, Germany). Furthermore, we used clear tape to remove spores from the plates and took pictures with the same microscope settings.

### Infection assays

For drop inoculations we cultured the 14 characterized isolates as described above, followed by a secondary incubation on fresh SNA plates for eight days under the same conditions. We harvested the spores by scraping them from the plates and placed them in water. Then we determined the spore concentrations under a microscope and diluted the solution to a concentration of 3×10^4^ spores per ml. We used five *Solanum* species for detached leaf infection assays: *S. chilense* (LA3111 and LA4117), *S. pennellii* (LA0716), *S. arcanum* (LA2133), *S. habrochaites* (LA0716) and *S. lycopersicum* (HEINZ1706). The plants originate from seed batches provided by the Tomato Genomics Resource Center (TGRC, Davis, CA) and the growth conditions in the greenhouse comprised 16h of light and a minimum temperature of 18°C. When harvesting the leaves, we chose several plants per population, collected leaves of approximately the same age and size, and randomized the leaves to minimize the effect of individual plants and leaf age. We placed the freshly cut leaves on wet tissue paper in boxes as in (Stam *et al*., 2017). To this end, we arranged rows with three leaves per species in a randomized order and added an additional leaf from each plant species for the positive control. These were inoculated with the *Alternaria solani* isolate 1117-1, which originates from *Solanum tuberosum* plants in Freising, Germany. This isolate reliably infects most potato and tomato cultivars (Nicole Metz, unpublished data). As negative control, we performed drop inoculations with sterile, distilled H_2_0. The smaller leaflets of *S. chilense* received one drop per leaflet and the bigger leaflets of *S. lycopersicum* received four drops per leaflet. The *Solanum* species *S. pennellii, S. arcanum* and *S. habrochaites* received two drops per leaflet. We placed the drops of 10 μl on the abaxial side of the leaves and stored the boxes at 25°C. After four days, we scored the infections (as described in Stam *et al*., 2017): When a drop did not result in symptoms or a small hypersensitive response lesion it counted as negative and when the spot displayed a large lesion or full proliferation of the pathogen it counted as positive. Results were calculated as the frequency of successful infections by dividing the number of successful infections by the number of inoculations. To obtain two biological replicates, we performed the whole setup twice.

## Results

### Fungal pathogens collected from wild tomato populations

In 2018 and 2019, we collected leaf samples from 81 different field locations (supp. table 4) spread over four regions ranging from northern Peru to northern Chile: Cajamarca, Lima, Tacna/ Arica and Antofagasta/San Pedro de Atacama (Fig. 1). Samples were taken from *Solanum arcanum, S. chilense, S. corneliomulleri, S. habrochaites, S. lycopersicoides, S. pennellii, S. peruvianum*, and S. *pimpinellifolium*. On nearly all sites at least some plants showed a combination of stress symptoms, ranging from very clear pathogen infection lesions (Fig. 2) to more generic browning or wilting of leaf tips. The severity of the symptoms varied between sites. In order to isolate *Alternaria-*like species, up to 10 symptomatic leaves from up to 5 different plants were collected on each site.

### Cultivation and molecular characterization

We were able to isolate a total of 372 fungal strains. Unfortunately, the success rate of the isolations was relatively low for the samples from the Cajamarca region, whereas for Antofagasta, the number of initially collected samples was already low. Nonetheless, we were able to successfully isolate fungal strains from each of the sampled regions and from all of the different hosts that occurred on the visited sites (Fig. 1 B-E). Visual inspection of all isolated cultures on plate showed that in the majority of cases we were able to isolate *Alternaria* or *Alternaria*-related specimens.

DNA isolation was successful for 211 isolates, from which 139 samples gave barcode sequences of *Alternaria* that were of sufficient quality to be included in the phylogenetic tree. The remaining 72 samples had to be removed because they had insufficient sequence quality, appeared to be unrelated species or possible contaminants.

### Species classification and phylogenetic analyses

To identify which species the samples belong to, we conducted a BLAST search of the sequences against the NCBI database and we followed a concatenation approach to construct a phylogeny. With the AB12PHYLO pipeline, we constructed a phylogenetic tree from all 139 samples with sufficient sequence quality and rooted it to *Stemphylium botryosum* (Fig. 3). It shows distinct groups of samples, with high support values for the nodes where the groups split. Inside the groups, the support values of some nodes are low because the samples are rather similar and their position would be interchangeable with their direct neighbors in the tree.

**Figure 3:**
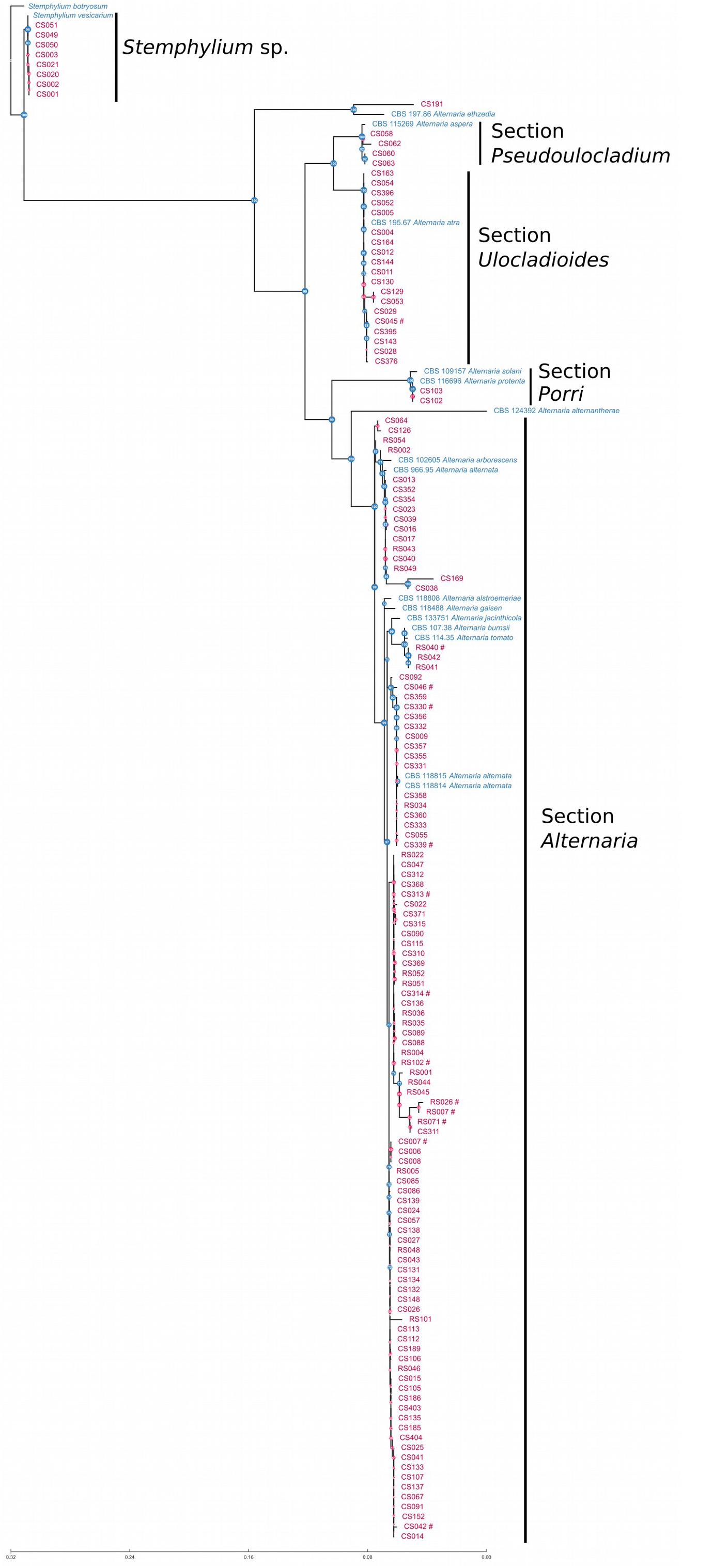
Phylogenetic tree from concatenated sequences The sequences of the four barcode markers Alt a 1, RPB2, TEF1 and ITS1F were concatenated for each sample and the reference sequences, then a phylogenetic tree was constructed from these sequences with RAXML-NG as part of the AB12PHYLO pipeline. References are labelled in blue while samples have dark red labels. Support of the clades is highlighted with color-coded node labels: blue nodes have a support value of more than 70 % or higher according to Transfer Bootstrap Expectations (TBE, Lemoine et al. 2008) and red nodes have 70 % or less support.

All eight samples with BLAST hits for *Stemphylium* and its sexual morph *Pleospora* group together with the references for this group. A single sample is close to the reference for *A. ethzedia* but the blast search of its barcode sequences gave hits for *A. ethzedia* and *A. rosae*, so the sample cannot be confidently determined to the species level. There are two groups of samples that belong to the section *Ulocladioides* and section *Pseudoulocladium*, respectively: 19 samples of *Alternaria atra* and four samples that group with *Alternaria aspera* but could not be identified to species level.

Two samples of large spored *Alternaria* were found. BLAST hits of their barcode markers include identifications as *A. protenta* and *A. solani*. In the tree, the samples group with both references, though they are slightly closer to *A. protenta* than to *A. solani*. Therefore, we can conclude that the samples belong to section *Porri* but identification as at species level remains less reliable.

The biggest group of samples with a total of 105 isolates belongs to the section *Alternaria*, which is sister to the reference for *A. alternantherae*. Inside this group, the samples cannot be assigned to species or species complexes because the resolution of the four barcode markers is not sufficient. This is demonstrated for example by the fact that the *A. arborescens* reference is grouping with one of the *A. alternata* references, not outside of one big clade representing *A. alternata*, and also by the fact that the references for *A. alstroemeriae, A. gaisen, A. jacinthicola, A. burnsii* and *A. tomato* would be expected to group outside of a branch with *A. arborescens* and *A. alternata* (compare Woudenberg *et al*., 2015). However, the BLAST hits suggest that most of the 105 isolates from this clade belong to the species *A. alternata*.

Additionally, we constructed phylogenetic trees for each individual barcode marker using the same settings in AB12PHYLO (supp. fig. S1). The individual trees demonstrate that phylogenies for the three barcode markers Alt a 1, RPB2 and ITS1F show very similar groups of samples, supporting the concatenated tree. The sequences for the barcode marker TEF1 are considerably shorter, so the phylogeny for this marker is less resolved and only delimits *Stemphylium* sp. from the other isolates. The placement of the reference for *Alternaria althernantherae* as the root of the small-spored section Alternaria is supported in the concatenated tree, the tree for Alt 1 a and the literature (Woudenberg *et al*., 2015), but not in the other single-marker trees. The tree for ITS1F shows a conflicting placement of sections *Ulocladioides, Pseudoulocladium* and *Porri* compared to the concatenated tree. Despite these inconsistencies, we show that the groups of isolates are consistent and reliably identified as members of the respective sections.

### Pathogen distribution and diversity

Most of the phylogenetic groups in the tree contained samples from different sampling regions and from different host plant species. The diversity of the phylogenetic groups in regard to their sampling regions and their host plants is illustrated in an alluvial plot (Fig. 4). For groups of pathogens with more than four isolates, we calculated diversity statistics in AB12PHYLO (Table 1).

**Figure 4:**
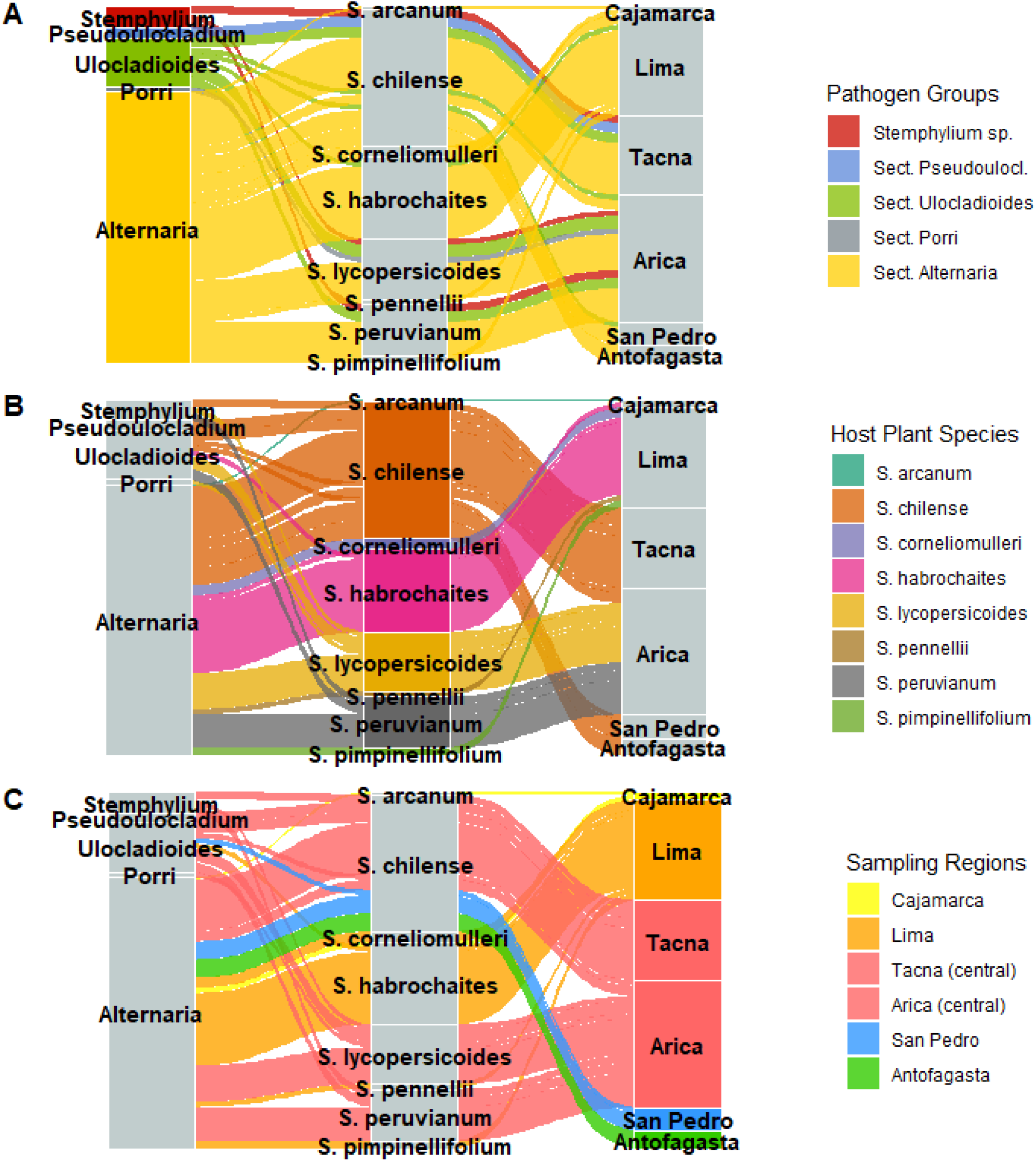
Alluvial plot illustrating the diversity of the samples A, B and C depict the same alluvial plot, each with different colors. This shows the diversity of the samples from the three perspectives of pathogen group as seen in the phylogenetic tree, host plant species and sampling region.

**Table 1:** Diversity statistics All diversity statistics were calculated with ab12phylo. Statistics are only shown for groups containing more than 4 isolates.

| Pathogen group | number<br>of<br>isolates | valid<br>sites | S<br>(segreg.<br>sites) | pi (nuc-<br>leotide<br>diversity) | Watterson's<br>Theta | Tajima's<br>D | unique<br>sequence<br>s |
| --- | --- | --- | --- | --- | --- | --- | --- |
| Stemphylium sp. | 8 | 995 | 0 | 0 | 0 | NA | 1 |
| Sect. Ulocladioides | 19 | 1256 | 4 | 0,000335 | 0,000911 | 1,080968 | 5 |
| Sect. Alternaria (small-spored) | 105 | 1515 | 123 | 0,007764 | 0,015534 | 1,646747 | 43 |
| considering all pathogen<br>species: |  |  |  |  |  |  |  |
| Lima | 39 | 1234 | 125 | 0,013331 | 0,023959 | 2,052769 | 21 |
| Tacna | 31 | 1492 | 284 | 0,050444 | 0,047647 | 4,090016 | 20 |
| Arica | 50 | 1259 | 266 | 0,050047 | 0,047169 | 3,842236 | 34 |
| San Pedro | 9 | 827 | 81 | 0,036947 | 0,036037 | 5,271321 | 4 |
| Antofagasta | 7 | 1438 | 12 | 0,003974 | 0,003406 | 6,348694 | 2 |
| *S. chilense* | 53 | 1528 | 327 | 0,043729 | 0,047158 | 3,342554 | 28 |
| *S. habrochaites* | 32 | 1450 | 151 | 0,015156 | 0,025858 | 2,233856 | 19 |
| *S. lycopersicoides* | 23 | 1232 | 219 | 0,050562 | 0,048163 | 4,24714 | 16 |
| *S. peruvianum* | 21 | 1036 | 201 | 0,054868 | 0,053927 | 4,181348 | 15 |
| considering only sect.<br>Alternaria: |  |  |  |  |  |  |  |
| Lima | 37 | 1238 | 60 | 0,07475 | 0,01161 | 2,33787 | 19 |
| Tacna | 20 | 1580 | 50 | 0,007045 | 0,00892 | 3,175919 | 10 |
| Arica | 31 | 1303 | 34 | 0,008787 | 0,006532 | 4,873817 | 18 |
| San Pedro | 7 | 829 | 29 | 0,010225 | 0,0114278 | 4,074119 | 3 |
| Antofagasta | 7 | 1438 | 12 | 0,003974 | 0,003406 | 6,348694 | 2 |
| *S. chilense* | 28 | 1513 | 80 | 0,007647 | 0,012585 | 2,223016 | 13 |
| *S. habrochaites* | 30 | 1470 | 64 | 0,007414 | 0,01099 | 2,539134 | 17 |
| *S. lycopersicoides* | 14 | 1323 | 31 | 0,00755 | 0,007368 | 4,40116 | 7 |
| *S. peruvianum* | 13 | 1081 | 29 | 0,010294 | 0,008645 | 5,209711 | 8 |

Upon closer inspection of the alluvial plot colored by the groups of pathogens as they can be seen in the phylogenetic tree, we see that most samples belong to section *Alternaria*, which was found on all host plant species in all sampling regions. This largest group is also the most diverse group, as there are for example more unique sequences per number of isolates in comparison to section *Ulocladioides* and the *Stemphylium* samples. When looking at the 105 isolates from the small-spored *Alternaria* section *Alternaria*, AB12PHYLO finds 43 unique sequences. The most common of these sequences occurs 21 times.

The other phylogenetic groups, especially section *Ulocladioides* and *Stemphylium* sp., exhibit less sequence diversity at the four investigated barcode sequences. The two isolates from section *Porri* are identical except one position in TEF1. The 19 isolates belonging to section *Ulocladioides* were collected at nine different locations in the regions of Lima, both central regions and in the southern mountain region near San Pedro de Atacama. Despite this large geographical spread, they only display five unique sequences: 15 of the 19 isolates share a unique sequence, the other four unique sequences occur only once. The four isolates grouping with *A. aspera* from section *Pseudoulocladium* were all collected from S. chilense at location S53 in Tacna. These four isolates have three unique sequences. The 8 *Stemphylium* isolates are from the central regions, but from 3 different locations with a different host species each. These 8 isolates have identical sequences within the alignment.

The alluvial plot colored by host plant species highlights which pathogens were found on the host plant species and in which regions the host plant species can be found. Host plants from which only few samples were collected only had pathogens belonging to the section *Alternaria*. When more samples could be taken from a host species, the pathogens were more diverse and belonged to different phylogenetic groups. *Solanum chilense* plants only grow in the central and southern sampling regions, but all dominant pathogen groups were found on it. The hosts *S. lycopersicoides* and *S. peruvianum* were only sampled in the central region Arica but also exhibited a variety of pathogen groups. The host species *S. habrochaites* from the northern regions mainly had pathogens from the section *Alternaria*.

These findings are reflected in the diversity statistics per species; when looking at all pathogens found on a given host species, *S. habrochaites* displays less diverse pathogens than *S. chilense, S. lycopersicoides* and *S. peruvianum*. We also calculated these diversity statistics for only small-spored *Alternaria*. The number of segregating sites (not corrected for sample size) is highest on *S. chilense* and *S. habrochaites* compared to *S. lycopersicoides* and *S. peruvianum*. The high values for pi hint that the small-spored pathogens on *S. peruvianum* are more diverse than on the other three host species.

Lastly, the plot colored by sampling region highlights that the central regions Tacna and Arica exhibited all kinds of *Alternaria* sections, while near Lima only pathogens from section *Alternaria* and a few specimens of *A. atra* were sampled. In both southern regions, fewer samples were collected, which all belong to section *Alternaria* except a few samples of *A. atra* in San Pedro de Atacama.

We find that both central regions Arica and Tacna harbor more diverse pathogens than the region around Lima, despite the fact that the samples were collected from 4 host plant species in Lima, but only 3 host plant species in Arica and only one host plant species, *Solanum chilense*, in Tacna. The climate and the diversity of the collected pathogens are very comparable in the two central regions Tacna and Arica, despite the different number of host plant species in these regions. Both southern regions seem to be slightly less diverse than the central regions. However, due to the much lower number of isolates collected in the southern regions, this might be a sampling artifact. The data is not sufficient to extrapolate whether collecting more isolates would increase the measured pathogen diversity.

### Morphological characterization of conidia from selected samples

Brightfield microscopy of the conidia from 14 samples shows characteristics congruent with the results of the molecular analysis. The removal of the conidia from the plate with sticky tape preserves the arrangement of the spores. All except one of the selected samples have conidia that grow in chains (supp. fig. S2), which is typical for section Alternata. Sample CS045 displays differences from the other samples. This is congruent with the molecular characterization: CS045 was identified molecularly as *A. atra* (section *Ulocladioides*).

When the spores are scraped from the plate and placed in water, the conidia are not arranged in chains and not attached to conidiophores anymore, but the septae of the conidia are visible with brightfield microscopy (Fig. 5). The shape of the conidia of CS045 is obovoid and they do not have beaks, which are the typical morphological characteristics of the section *Ulocladioides* (Woudenberg *et al*., 2013; Lawrence *et al*., 2016). The other samples belong to the section *Alternaria* according to our phylogenetic analysis and in fact display morphological characteristics of this section (as described in Woudenberg *et al*., 2013; Lawrence *et al*., 2016).

**Figure 5:**
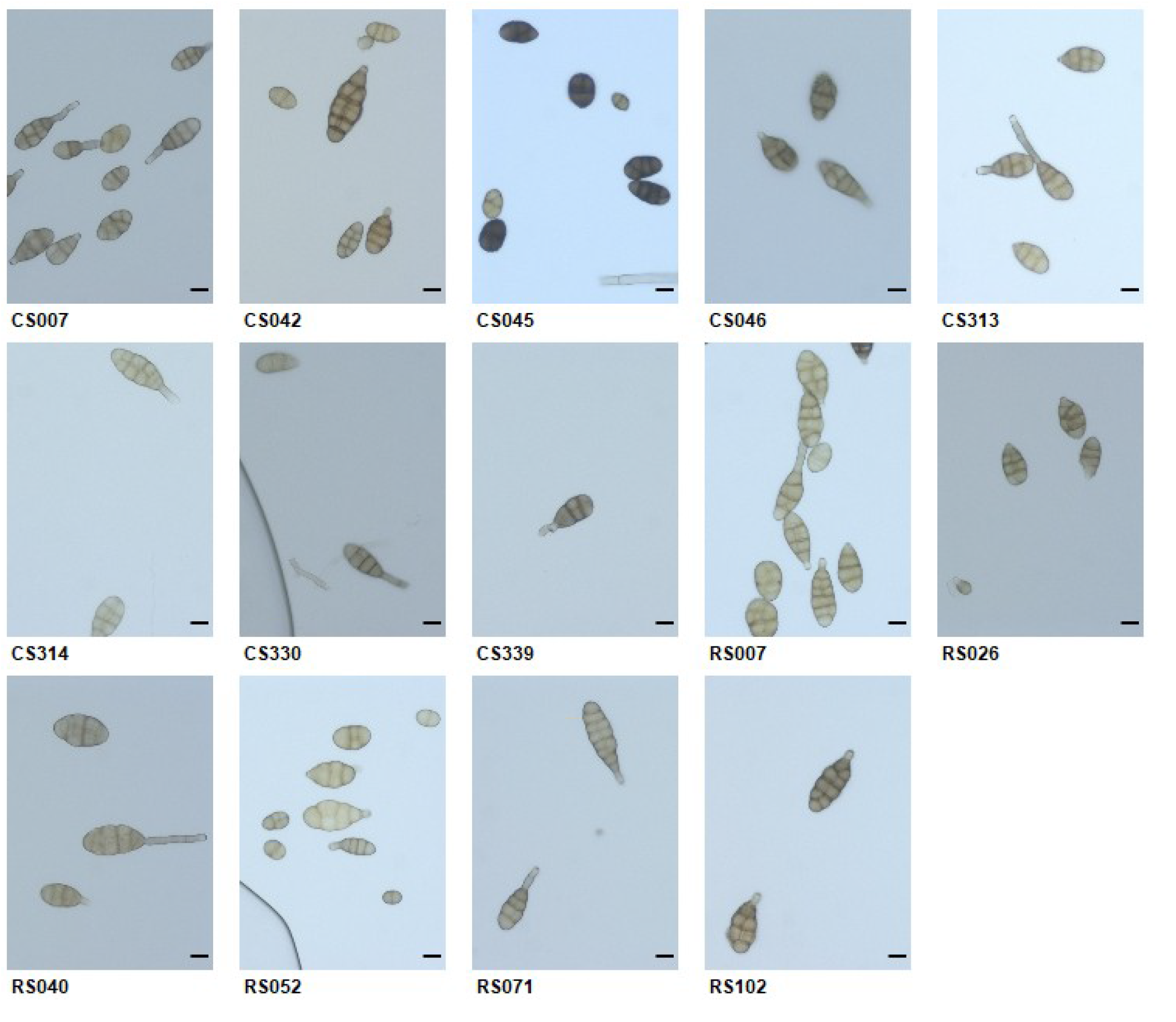
Conidia in water All isolates grew on SNA plates for eight days. They were scraped from the plates and placed in a drop of water for microscopy. Scale bars = 10 μm

The conidia are small or moderate in size, their form is obclavate or long ellipsoid, and they are septate with slight constrictions near some septa, in most cases with a few longitudinal septa. Interestingly, the two samples CS330 and CS339 grow slightly slower on SNA plates and exhibit very few and only juvenile conidia after growing on SNA plates for eight days, while all other samples produced plenty of mature spores during this time. In conclusion, the morphological characteristics of the selected samples are congruent with the results of the molecular characterization.

### Infection assays

To confirm that the collected isolates are pathogens on wild tomato species we conducted a detached leaf infection assay. To this end, we infected leaves of four different wild tomato species and one cultivated tomato species with drops containing spores of the fourteen above-mentioned isolates. The infectivity of the isolates on the different tomato species varied depending on the isolate as well as the tomato species (Fig. 6). Our positive control, an *A. solani* isolate from Germany, infected all tomato species including the cultivated tomato cultivar HEINZ1706. Differences between the plant species’ reactions to the *A. solani* isolate can be observed and indicate that *S. pennellii* was most resistant and *S. chilense* most susceptible among the wild tomato species. The isolates collected from wild tomato plants in Peru and Chile only rarely caused infections on the cultivated tomato, possibly indicating a certain degree of host specificity. The wild tomato species *S. chilense* generally shows the highest infection rates, only four of the fifteen isolates caused more infections on another tomato species than on *S. chilense*.

**Figure 6:**
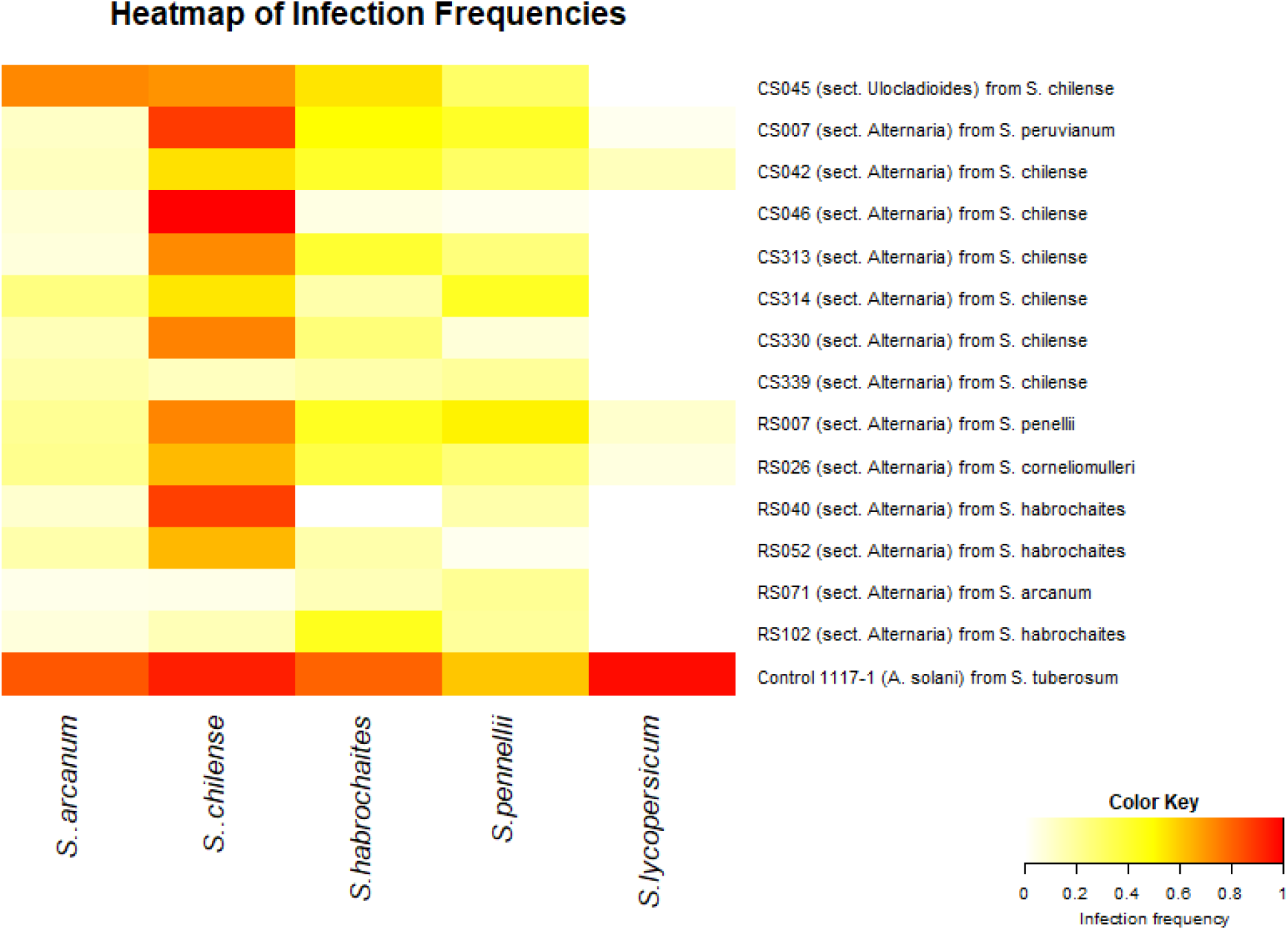
Heatmap visualizing the infection frequencies Drop inoculations on detached leaves showed that all tested isolates have the ability to infect wild tomato species. The successful infection frequency varied depending on the isolate as well as the tomato species. The rows show the isolates with sampling number, affiliation to species or group in the multigene phylogeny and original host plant. Each column represents a tomato species: We tested four wild tomato species and the cultivated tomato *S. lycopersicum*. Infection frequencies range from 0 (0 % of the inoculation drops resulted in an infection) to 1 (100 % of the inoculation drops resulted in an infection).

## Discussion

Understanding resistance mechanisms in wild crop relatives is an important pillar of modern plant breeding. To place such findings into context, information of naturally occurring pathogens is needed. Here we present first insights into the diversity of *Alternaria* on wild tomato species in their natural habitat. Our study benefits from a very broad sampling design. This allows us to capture the diversity from different regions in two countries. The largest distance between sampling sites is more than 2000 km. Naturally, not all *Solanum* host plant species occur in all climates and regions, but with up to 4 host species per region and a total of 8 host species, the sampling design facilitates insights into the effect of the wild tomato species.

### Complexities with the classification of *Alternaria* species

The phylogenetic tree of concatenated samples allows us to assign nearly all 139 isolates to a section with high node support. Most of these classifications are supported by the phylogenetic trees of separate barcode markers. The markers Alt 1 a and RPB2 are most specific for *Alternaria*-like species among our four markers. The tree for ITS1F also shows relatively similar clades, although this marker is rather conserved in *Alternaria*-related fungi (e.g. Woudenberg *et al*., 2015; Dettman and Eggertson, 2021). Due to technical difficulties, the sequences for TEF1 were far shorter than expected (61 aligned characters in our study compared to 241, 201 and 240 aligned characters in (Woudenberg *et al*., 2015; Landschoot *et al*., 2017; Ding *et al*., 2019), respectively.)

In our study, the resolution of the markers to distinguish between species within the small-spored section *Alternaria* was lower than previously reported in literature. According to Woudenberg et al. (2015) and Landschoot et al. (2017), *A. alternata* can be differentiated from the *A. arborescens* species complex with the Alt 1 a and TEF1 markers. The RPB2 marker should also distinguish between *A. alternata* and other species. However, our phylogenetic tree shows that only approximately half of our small-spored isolates cluster with the references for *A. alternata*. Furthermore, one of the references for *A. alternata* clusters closely together with *A. arborescens* while the other references for *A. alternata* can be found in a sister clade. The other half of our small-spored isolates does not group together with any known reference sequence. This might indicate that the small-spored *Alternaria* species infecting wild *Solanum* are even more diverse than the *Alternaria* species described so far. All in all, we have to conclude that the resolution of the four employed barcode markers is insufficient to distinguish between closely related species. This is in line with a recent study by Dettman and Eggertson 2021, who state that the markers that we employed are capable of placing an isolate into a section, but that section-specific markers would be necessary for a better resolution within section *Alternaria* (Dettman and Eggertson, 2021). They demonstrate that the existing genetic markers are located in regions of the genome with low gene/species concordance and propose candidate genes which would provide better resolution, which could be used in future studies. However, only whole genome sequences of several lineages in *Alternaria alternata* could reveal whether *A. alternata* forms a big panmictic population with cryptic sex (Meng *et al*., 2015) or has diversified into genetically distinct lineages that could be considered as independent species.

In this context it is not surprising that many species related to *Alternaria* and especially members of the small-spored section *Alternaria* have received several taxonomic revisions. Morphological characteristics support our molecular classification on section level, but do not provide identification to species level. Such characteristics overlap between species, depend on growth conditions, and may not reflect the evolutionary relationship between species within the sections (Dettman and Eggertson, 2021). Especially the conidial morphology is variable and depends on environmental factors, which has already lead to wrong species classifications in the past (Thomma, 2003).

Due to the low resolution within the small-spored section, we can only assume that the majority of isolates in the small-spored clade should be considered *A. alternata* sensu stricto. The whole section *Alternaria* is often referred to as “*alternata* clade” (Dettman and Eggertson, 2021), so in a broader sense the small spored isolates might be referred to as *A. alternata*.

### Prevalence of small-spored isolates

The majority of our collected isolates are small-spored and belong to *Alternaria* section *Alternaria*. The fact that only two isolates belong to section *Porri* is surprising, because large spored species like *A. linariae* are reported as major problems on tomato (e.g. Peixoto *et al*., 2021). According to Adhikari et al. 2020, *A. alternata* on tomato is neglected and poorly understood in comparison to *A. linariae* (Adhikari *et al*., 2020). However, there are more and more reports of small spored *Alternaria* being dominant on tomato crops (e.g. (Bessadat *et al*., 2017; El Gobashy *et al*., 2018; Kokaeva *et al*., 2018). Seeing that 75% of our isolates belong to section *Alternaria*, we report that small spored species are dominant on wild tomato as well.

The higher prevalence of small-spored pathogens might be caused by seasonality. Adhikari et al. 2020 state that they collected their *A. alternata* isolates later in the season than their *A. solani* isolates (Adhikari *et al*., 2020). Both our sampling trips took place in early March, which is late summer on the southern hemisphere and should allow for the detection of large-spored isolates. However, cultivated crops have very clear growing seasons, while the host plants in this study are perennial, so the seasons would affect the abiotic conditions more than the host plant. While the first description of *A. alternata* on potato reported that the disease development increases during the season similar to *A. solani* (Droby *et al*., 1984), Vandecasteele et al. 2018 found that small-spored species are predominant throughout the season (Vandecasteele *et al*., 2018). We conclude that we should have detected large-spored isolates if they had been omnipresent on wild tomato plants but recommend further sampling trips in different seasons to capture an even greater diversity of *Alternaria*-like specimens.

Most wild tomato sampling sites were rather removed from crops, making the studied pathosystem a truly wild system in most cases. However, some of the sampling sites are in proximity to potato crops (areas with higher elevation) and tomato crops (areas with lower elevation). Local farmers on at least one of the potato cultivations reported problems with *A. solani*. The fact that hardly any *A. solani* was found on the wild tomatoes might indicate that there is no cross-contamination from crops to the wild. Peixoto et al. 2021 found several *Alternaria* pathogens from different solanaceous plants in Brazil to cause early blight symptoms on tomato. As these had originally been collected from persistent weeds, they claim it is likely that these plants can act as alternative hosts and might become a source of inoculum to important crops (Peixoto *et al*., 2021). Interestingly, our collected pathogens from wild tomato plants did not infect cultivated tomato in an infection assay. We therefore assume that there is neither cross-contamination from the wild tomato plants to the field nor vice versa.

### Diversity of the collected pathogens

Taking into account minor differences in definitions of genotype and unique sequence, we find more unique sequences than a study investigating *A. alternata* diversity from potato crops in Wisconsin. Ding et al. 2019 collected *A. alternata* and *A. solani*. With five barcoding markers, three of which we also employed, they grouped their 40 *A. alternata* isolates in five genotypes (Ding *et al*., 2019). The 105 small-spored isolates in our study showed 43 unique sequences and formed approximately 4 or 5 larger groups in the concatenated phylogenetic tree, which consist of a dominant sequence but also several very similar sequences. The higher number of unique sequences in our study is expected, because our sampling design includes several host plants in many geographical locations from different climatic regions, while Ding et al. 2019 only sampled cultivated potato three regions (approximately 30 km apart).

The most common genotype in Ding et al. 2019 represented 58 % of their 40 *A. alternata* isolates. The most common unique sequence in our study occurred in 20% of our small-spored isolates (21 of 105 isolates). The observed diversity in our data thus points towards a larger genetic biodiversity in the wild pathosystem, which will be further investigated with a whole genome study in the future.

Ding et al. 2019 found a mixture of genotypes at each location and retrieved the genotypes in the same relative abundances five years after their first collection. Furthermore, they argue that *A. alternata* are genetically mixing because they did not find a difference in virulence between *A. alternata* isolates from different regions and no distribution pattern of genotypes. We also found more than one unique sequence on average at each collection location and found that all isolates infect all hosts. Our two collection trips took place in different regions, so we cannot compare different time points, but we find that groups of unique sequences encompass isolates from both trips, indicating that samples with this sequence have been collected in both years. Adhikari et al. 2020 collected *A. alternata* from tomato crops in Stokes county, North Carolina, in 2012 and 2014. They defined haplotypes based on the GPDH sequence and report that the same haplotype was dominant in both years. This most common haplotype was also found in the other three counties they sampled. In fact, most of the haplotypes they defined occur in several of their sampling locations (Adhikari *et al*., 2020). All three studies therefore point towards a broad distribution of small-spored *Alternaria* haplotypes over space and time.

Small-spored *Alternaria* exhibit greater sequence diversity than the other pathogen sections. Ding et al. 2018 find that *A. alternata* isolates were more diverse than *A. solani* isolates (Ding *et al*., 2019). Adhikari et al. 2020 report *A. alternata* to show more sequence diversity, especially higher values for nucleotide diversity pi and Watterson’s theta, than *A. linariae* or *A. solani* (Adhikari *et al*., 2020). We did not collect enough large-spored specimens to draw meaningful conclusions about this group, but we also find that the small-spored section *Alternaria* shows more sequence diversity than the section *Ulocladioides* and the *Stemphylium* sequences. Adhikari et al. 2020 collected their *A. alternata* isolates later in the season than their large-spored isolates, so the high diversity of *A. alternata* is unlikely to be an artifact from sampling during a season with mostly *A. alternata*.

The high diversity within section *Alternaria* might be caused by recombination (Ding *et al*., 2019). Despite the fact that *Alternaria* reproduces asexually, there are strong indications that recombination occurs within *A. alternata* (Meng *et al*., 2015).

### Adaptation to host plant species and habitat

As not all host plant species occur in all regions, it is impossible to completely disentangle the adaptation to the host and to the climate. The prevalence of *Alternaria* disease on potato depends on several environmental factors. When wet and dry periods alternate, the growth of *Alternaria* hyphae is favored, while periods with heavy rain favor sporulation. Infections are favored under humid conditions and temperatures of 24-29° C. These effects often seem contradictory, for example when higher disease severity is observed at lower temperatures due to optimal wetness-related conditions (Vandecasteele *et al*., 2018). Theoretically, we would therefore expect the highest pathogen pressure in the northern sampling regions. Despite the very contrasting climatic conditions of our sampling regions, we find very broad distributions of the collected *Alternaria* pathogens. Using only barcode markers, we could not find any signs of climate adaptation.

In other wild pathosystems, the pathogens are usually most aggressive on their original host (e.g. Tack *et al*., 2012). In our experiments, we did not find such results. However, it should be noted that the tomato accessions used in this experiment were randomly selected to represent a host species and do not stem from the original locations where the pathogens were sampled. For example, the seed stock used for *S. pennellii* originates from the region Arequipa in Peru, where no sampling was conducted.

As we rarely found wild specimens of large-spored species which are dominant on potato and tomato crops, and the wild pathogens did not infect cultivated tomato in our infection assay, we conclude that the collected wild pathogens are likely adapted to their wild host plants. Besides this, we did not find any signs of host specificity using the barcode sequences. From the genetic data gathered in this study, we see a broad distribution of the pathogen groups, and no pathogen group was specific to a host species or a region. These findings are congruent with the study on cultivated tomato in North Carolina. Adhikari et al. 2020 report that they do not find any association between haplotype or species and host geographic location (Adhikari *et al*., 2020). Weir et al. 1998 showed a host specialization of *A. solani* by determining that the genetic distance between isolates from tomato and potato hosts is significantly large (Weir *et al*., 1998). Interestingly, they do not report this for their *A. alternata* isolates from tomato and potato, though the difference between solanaceous hosts and citrus was clearly visible in their RAPD study.

The ability of *Alternaria* to infect a certain host often depends on the ability to produce host-specific toxins (HST). HST are chemically diverse and have different sites of action, but all trigger cell death. Biosynthesis genes for HST usually cluster together on a supernumerary (sometimes called accessory) chromosome in *A. alternata* pathotypes. It is hypothesized that Alternaria became pathogenic by acquiring the HST genes through horizontal gene transfer (HGT) of this supernumerary chromosome (Thomma, 2003). Therefore, we will need to employ whole genome sequencing data to elucidate host specificity. As with climate adaptation, we conclude that we cannot find signs of host adaptation using only barcode markers. Hopefully, a study of whole genomes will elucidate the genomic basis of adaptation in the future.

## Summary

*Alternaria* spp. and related fungi are common pathogens on wild tomato plants. With an exceptionally broad sampling design, we show that these pathogens occur on all eight sampled host species in six regions of Chile and Peru, covering diverse climatic conditions and more than 2000 km geographical distance. Sequencing genetic barcode markers showed that predominantly small-spored species of *Alternaria* of the section *Alternaria*, like *A. alternata*, caused infections, but also by *Stemphylium* spp., *Alternaria* spp. from section *Ulocladioides*, and related species. Morphological observations and an infection assay confirmed the molecular analyses. Comparative genetic diversity analyses show a larger diversity in this wild system than in studies of cultivated *Solanum* species.

As the wild pathogens did not infect the cultivated tomato in the infection assay and hardly any large-spored *Alternaria* species were collected from wild plants despite being prominent on solanaceous crops in the viscinity of some sampling locations, we conclude that there is no cross-contamination between the wild system and cultivated crops. However, this study is not only relevant to scientists studying wild systems but also to crop protection and plant breeding efforts, especially since *A. alternata* has been reported as growing problem on tomato crops.

## Acknowledgments

We thank Regina Dittebrand for help with fungal propagation and Leo Kaindl for constant improvements to the AB12PHYLO software. This work was supported by DFG grants 1547/2-1 and 1547/4-1.

## Supplementary Materials

Supplementary figure S1: Phylogenetic trees of barcode markers

Supplementary figure S2: Conidia on sticky tape

Supplementary table S1: Primers used for barcode sequencing

Supplementary table S2: PCR conditions

Supplementary table S3: Accession numbers of reference sequences

Supplementary table S4: Sampling sites

Supplementary table S5: List of isolates

